# Placenta microbiome diversity is associated with maternal pre-pregnancy obesity and placenta biogeography

**DOI:** 10.1101/659797

**Authors:** Paula A. Benny, Fadhl M. Al-Akwaa, Corbin Dirkx, Ryan J. Schlueter, Thomas K. Wolfgruber, Ingrid Y. Chern, Suzie Hoops, Dan Knights, Lana X. Garmire

## Abstract

Recently there has been considerable debate in the scientific community regarding the placenta as the host of a unique microbiome. No studies have addressed the associations of clinical conditions such as maternal obesity, or localizations on the placental microbiome. We examined the placental microbiome in a multi-ethnic maternal pre-pregnant obesity cohort using controls for environmental contaminants and an optimized microbiome protocol to enrich low bacterial biomass samples. We confirmed that a distinct placenta microbiome does exist, as compared to the environmental background. The placenta microbiome consists predominantly of *Lactobacillus, Enterococcus* and *Chryseobacterium*. Moreover, the microbiome in the placentas of obese pre-pregnant mothers are less diverse when compared to those of mothers of normal pre-pregnancy weight. Lastly, microbiome richness also decreases from the maternal side to fetal side. In summary, our study reveals associations of placental microbiome with placenta biogeography and with maternal pre-pregnant obesity.

## Introduction

The human microbiome is the collection of microorganisms which reside on or in human organ systems. Symbiosis is the mutually beneficial relationship between humans and these microorganisms whereas dysbiosis is the imbalance of the human microbiome. Dysbiosis has recently been associated with diseases and abnormalities(1), including pre-term birth. In particular, subjects who experienced pre-term labor had lesser *Lactobacillus* in their microbiome as compared to term gestation subjects. In addition, wider bacterial diversity was noted in pre-term pregnancies as compared to controls, including those associated with the vaginal microbiome such as Ureaplasma species and those associated with the oral microbiome such as Streptococcus thermophilus (2-4).

While microbiome existence has been widely recognized in human organs such as the skin, gut and vagina, there have been debates on the presence of a distinct microbiome in placental tissue (2-8). The landmark report by Aagaard et al (5) demonstrated the presence of a unique placental microbiome. Subsequently, other studies have attempted to characterize the placental microbiome as well (7, 9). On the contrary, some other studies (6, 10) refuted this idea. Using 16S rRNA sequencing approach, the authors argued that the placental microbiome reported in the Aagaard study was due to environmental or reagent contamination, which were not accounted for in the original study through the inclusion of adequate controls for contamination.

In light of the debate above, we embarked on a new study on a group of women undergoing elected caesarean sections in a sterile environment, to eliminate other potential sources of bacterial contamination associated with vaginal births. Additionally, to determine the possible association between maternal obesity status and placental microbiome, we divided them into cases and controls according to their pre-pregnancy weight: either pre-pregnant obese (BMI>30) or normal-weighted (18.5<BMI<25). Furthermore, we collected multiple placenta samples per patient, from the maternal, fetal and intermediate layers, along with various environmental controls including delivery room airswabs, laboratory airswabs and unopened reagents. To account for the low bacterial biomass, we developed an optimized protocol to enrich the V4 region of bacterial 16S rRNA genes, and for controlling for environmental contaminants. The results suggest that there indeed exists a placental microbiome even after controlling for contamination. The placental microbiome between pre-pregnant obese and normal pre-pregnant weight women was markedly different, in that the former group has a less diverse microbiome.

## Materials and Methods

### Sample collection

Placenta samples were collected from pregnant mothers admitted for elected full-term cesarean section at ≥ 37 weeks gestation at Kapiolani Medical Center for Women and Children, Honolulu, HI from November 2016 through September 2017. Such procedure minimized introduction of other bacteria associated with vaginal births as well as bacterial contamination from air during births. The study was approved by the Western IRB board (WIRB Protocol 20151223). Women with preterm rupture of membranes (PROM), labor, multiple gestations, pre-gestational diabetes, hypertensive disorders, cigarette smokers, HIV, HBV, and chronic drug users were excluded from the study. Patients meeting inclusion criteria were identified from pre-admission medical records with pre-pregnancy BMI ≥30.0 (obese) or 18.5-25.0 (normal pre-pregnancy weights). Demographic and clinical characteristics were recorded, including maternal and paternal ages, maternal and paternal ethinicities, mother’s pre-pregnancy BMI, pregnancy net weight gain, gestational age, parity, gravidity and ethnicity. Placenta samples were obtained equally distant from the cord insertion site and the placenta edge. Placenta samples were isolated (0.5cm^3^) from the maternal, fetal and intermediate areas using sterile surgicals. To consider all possible sources of environmental contaminations, air swab samples were obtained by waving the airswab in the air in the surgery room, the pathology lab where the placenta biopsies were collected, and the research laboratory where extraction was carried out. Unopened airswabs were also used as a control.

### Extraction of genetic material

MOBIO Powersoil DNA Kit (#12888-50) was used to extract DNA from placenta samples. 300mg of placenta was homogenized, heated for 65°C and vortexed in a horizontal bead beater for 10 minutes. DNA was extracted from lysates by putting them through the MOBIO kit following the manufacturer’s protocol. Extracted DNA was quantified and QC-checked using Nanodrop.

### Bacterial DNA enrichment

Given the very low bacterial mass, an enrichment step was performed to remove host DNA contamination and improve 16S specific amplification. (NEBNext Microbiome DNA Enrichment Kit, # E2612L). Samples were enriched in sets of 8 for optimal enrichment of bacterial DNA. DNAs were incubated with NEBNext magnetic beads for 15 minutes. Beads containing human host DNA were precipitated using a magnet, leaving microbial DNA in the supernatant.

### qPCR amplification

qPCR was performed to determine 16S counts within extracted samples. Isolated microbial DNA was amplified using primers to the hypervariable V4 region of 16S rRNA gene, similar to others (7, 11). Forward primer – TCGTCGGCAGCGTCAGATGTGTATAA GAGACAGGTGCCAGCMGCCGCGGTAA. Reverse primer – GTCTCGTGGGCTCG GAGATGTGTATAAGAGACAGGGACTACHVG GGTWTCTAAT. PCR was performed using KAPA HiFidelity Hot Start Polymerase; 95°C for 5 mins, 98°C for 20s, 55°C for 15s, 72°C for 1 minute for 25 cycles, 72°C for 5 minutes. After the 25 cycles of amplification, V4 specific amplicons were observed by 2% agarose gels and Agilent Bioanalyzer traces. V4 amplicon was detected at the expected size of 290bp. Samples were pooled, size-selected and denatured with NaOH, diluted to 8pM in Illumina’s HT1 buffer, spiked with 15% PhiX, and heat denatured at 96°C for 2 minutes immediately prior to loading. A MiSeq600 cycle v3 kit was used to sequence the samples, following the manufacture’s protocol.

### Bioinformatics analysis

The 16S rRNA gene reads were analyzed using the pipeline shown in Supplementary Figure 1. Reads were stitched using Pandaseq (12) using 150bp and 350 bp as the minimum and maximum lengths of the assembled reads respectively. Operational taxonomic units (OTUs) were created by clustering the reads at 97 % identity using UCLUST (13). Representative sequences from each OTU were aligned using PyNAST (14), and a phylogenetic tree was inferred using FastTree v. 2.1.3 (15) after applying the standard lane mask for 16S rRNA gene sequences, Pairwise UniFrac distances were computed using QIIME (16). Permutation tests of distance and principal coordinates analyses were performed using the MicrobiomeAnalyst, a web-based tool for comprehensive exploratory analysis of microbiome data (17). Taxonomic assignments were generated by the UCLUST consensus method of QIIME 1.9, using the GreenGenes 16S rRNA gene database v. 13_8 (18). We used Phyloseq R package to compute alpha and beta diversity(19). We used SourceTracker (version 1.0.1) to estimate the percentage of OTUs in placental samples whose origin could be explained by their distribution in the airswabs (20).

## Results

### Demographic and clinical characteristics of the cohort

Our cohort consisted of 44 women from three ethnic groups including Caucasians, Asians and Native Hawaiians, who underwent scheduled full-term caesarean deliveries in Kapiolani Medical Center for Women and Children, Honolulu, Hawaii from November 2016 through September 2017. The patients were included based on the inclusion and exclusion criteria described earlier (Methods section). In order to test if there is microbiome difference associated with maternal pre-pregnancy obesity, the subjects were recruited in two groups: normal pre-pregnant weight (18.5<BMI<25) and pre-pregnant obese (BMI> 30) group. The patient demographical and clinical characteristics are summarized in Table 1. Maternal ages, gestational weight gain and gestational age differences between the cases and controls are not statistically significant, excluding the possibility of significant confounding from these factors. Maternal pre-pregnant obesity, however, is associated with increasing parity and gravidity (P<0.05). The variation in recruited cases versus controls in each ethnic background reflects the multi-ethnic population demographics in Hawaii.

**Table 1:**
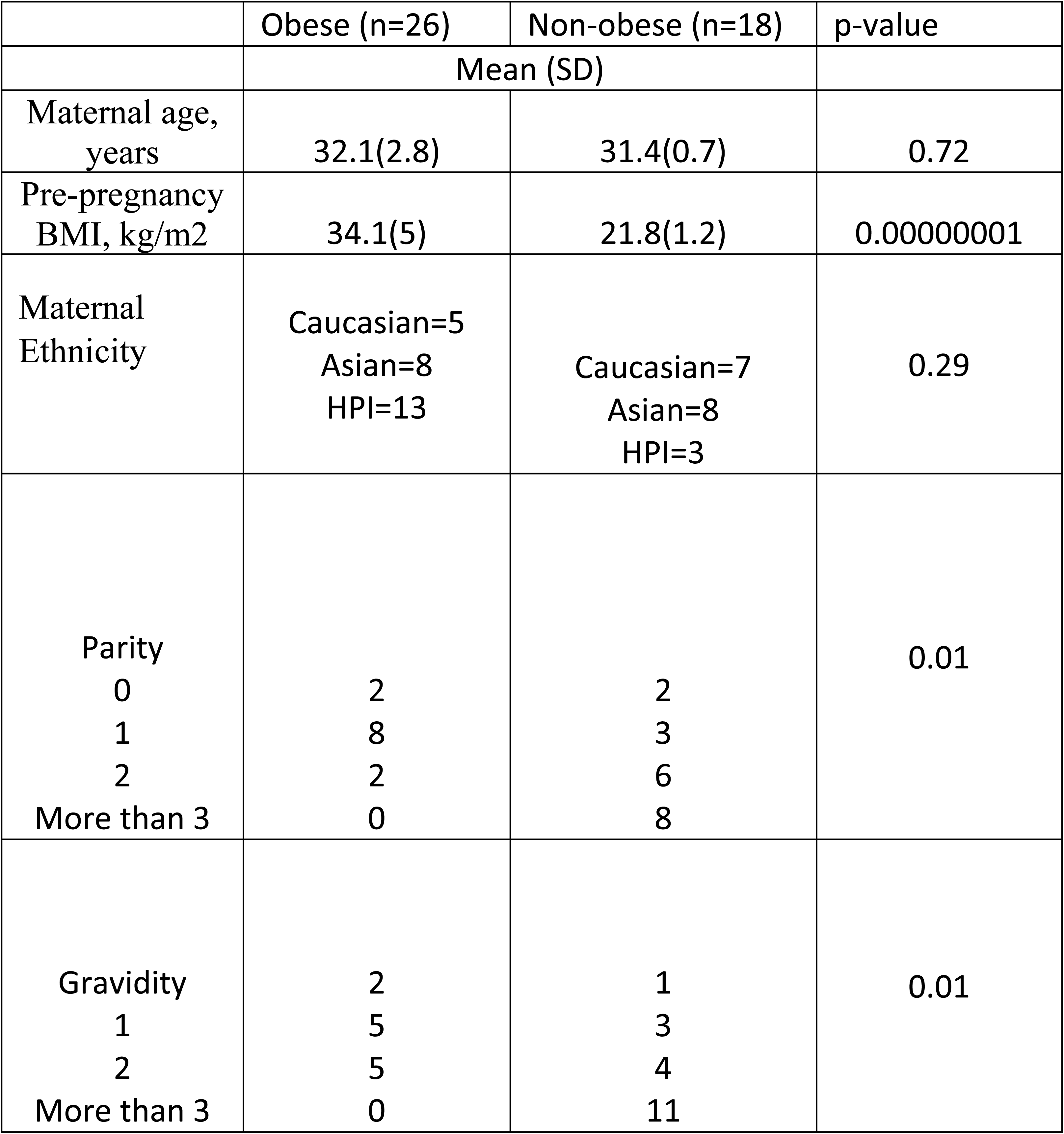
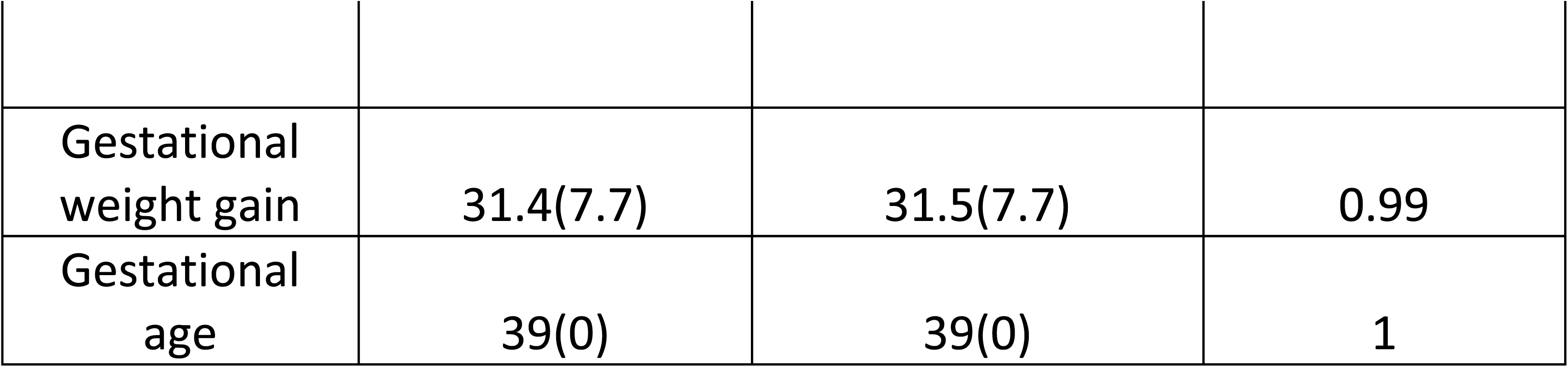
Clinical characteristics of the cohort.

### Enrichment of the placental microbiome

We first performed qPCR to determine the 16S rRNA copy numbers within extracted samples. As shown in Figure 1A, placenta samples contain significantly more copies of 16S as compared to airswab or water controls; placenta 73,595+/-1485 mol/µl, airswab 83 +/-43 mol/µl, water 24 +/− 11 mol/µl. The difference of 16S transcript numbers between placentas and airswab/water is significant (P<0.05). Given the extremely low bacterial biomass in placenta, we implemented an enrichment step in bacterial V4 region to remove human DNAs (Methods). As showing in Figure 1B, unenriched V4 samples yield much lower total reads (median: 68,468) as compared to enriched V4 samples (median: 516,479), suggesting the success of the experimental protocol. Furthermore, V4 amplicons post-PCR on the agarose gel show the specific band of 290bp – the expected size of V4 amplicons, confirming successful 16S amplification of placenta samples (Figure 1E). This band is missing in the negative controls of airswabs and water, confirming that the bacterial DNAs are below detectable levels in the environmental controls.

**Figure 1:**
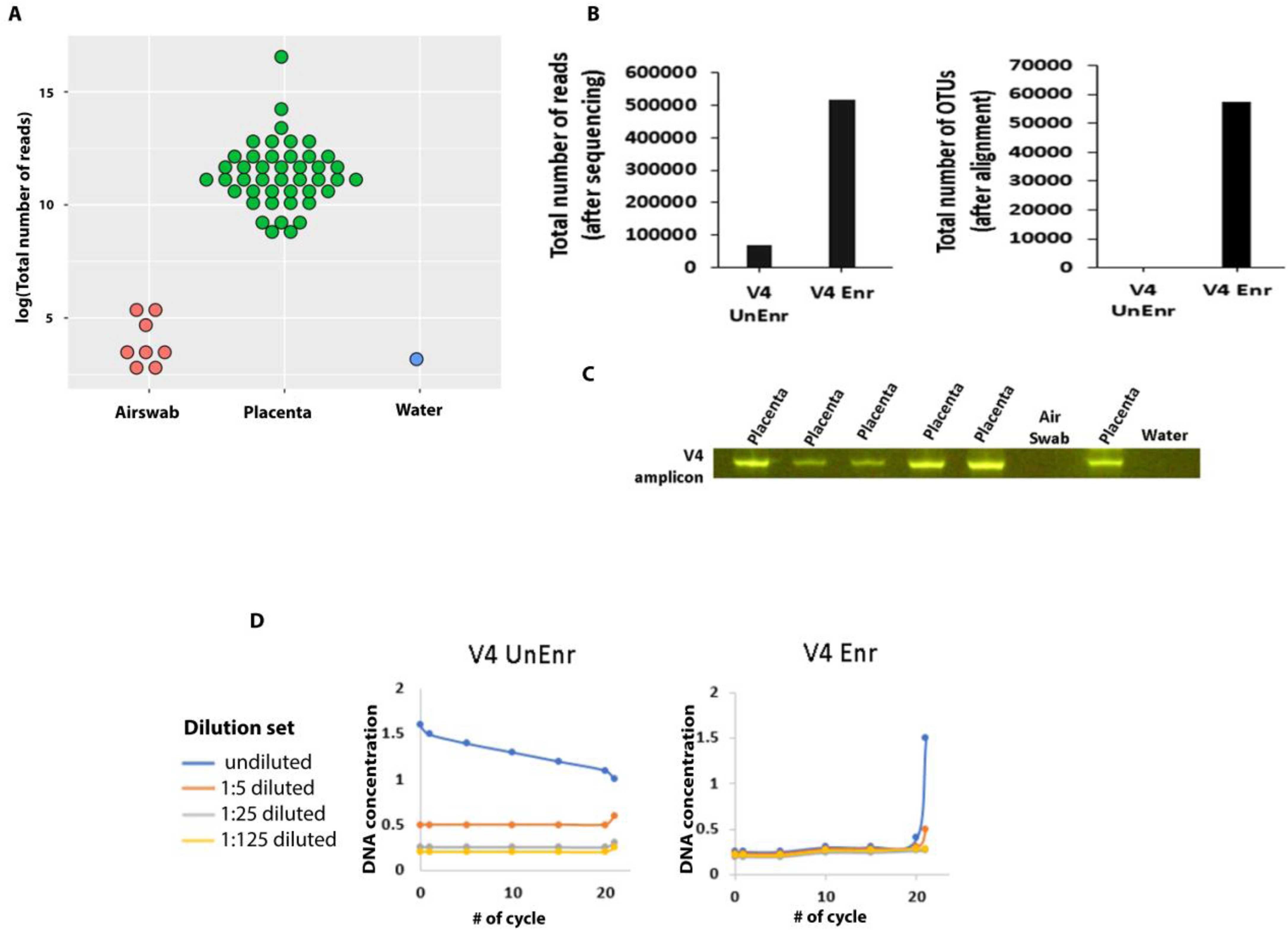
(A) 16S counts of placenta and controls (airswabs and water) using qPCR. (B) Total number of reads in enriched and unenriched samples while varying primer types. (C) Total number of OTUs (after alignment to Greengenes database) in enriched and unenriched samples. (D) qPCR analysis to justify the need for a microbial enrichment step and using V4 primers during the amplification step. (x-axes is the number of cycle, y-axes is the DNA concentration and lines are different DNA dilution, undiluted (blue line), a 1:5 template (orange line), a 1:25 template (grey line) and a 1:125 template (yellow)) (E) Agarose gel run showing specific V4 amplicon (290bp) detected in placenta samples, which are not present in airswab or water controls.

### Placenta microbiome is different from the environmental background

We implemented a bioinformatics analysis workflow as shown in Figure S1. We aligned the 16S sequencing reads using Greengenes database. The enriched samples using V4 primers detect on average 57,468+/-2,859 operational taxonomic units (OTUs), compared to 233+/-36 OTUs on average from un-enriched samples (Figure 1C), again highlighting the strength of the enrichment step following DNA extraction. Furthermore, the principal coordinates analysis (PCoA) plot based on OTUs shows that indeed airswabs and placental microbiome are distinct and separable into two clusters (Figure 2B). Alpha diversity was higher in placenta than airswabs (t-test, p = 6.4947e-07) (Figure 2A).

**Figure 2:**
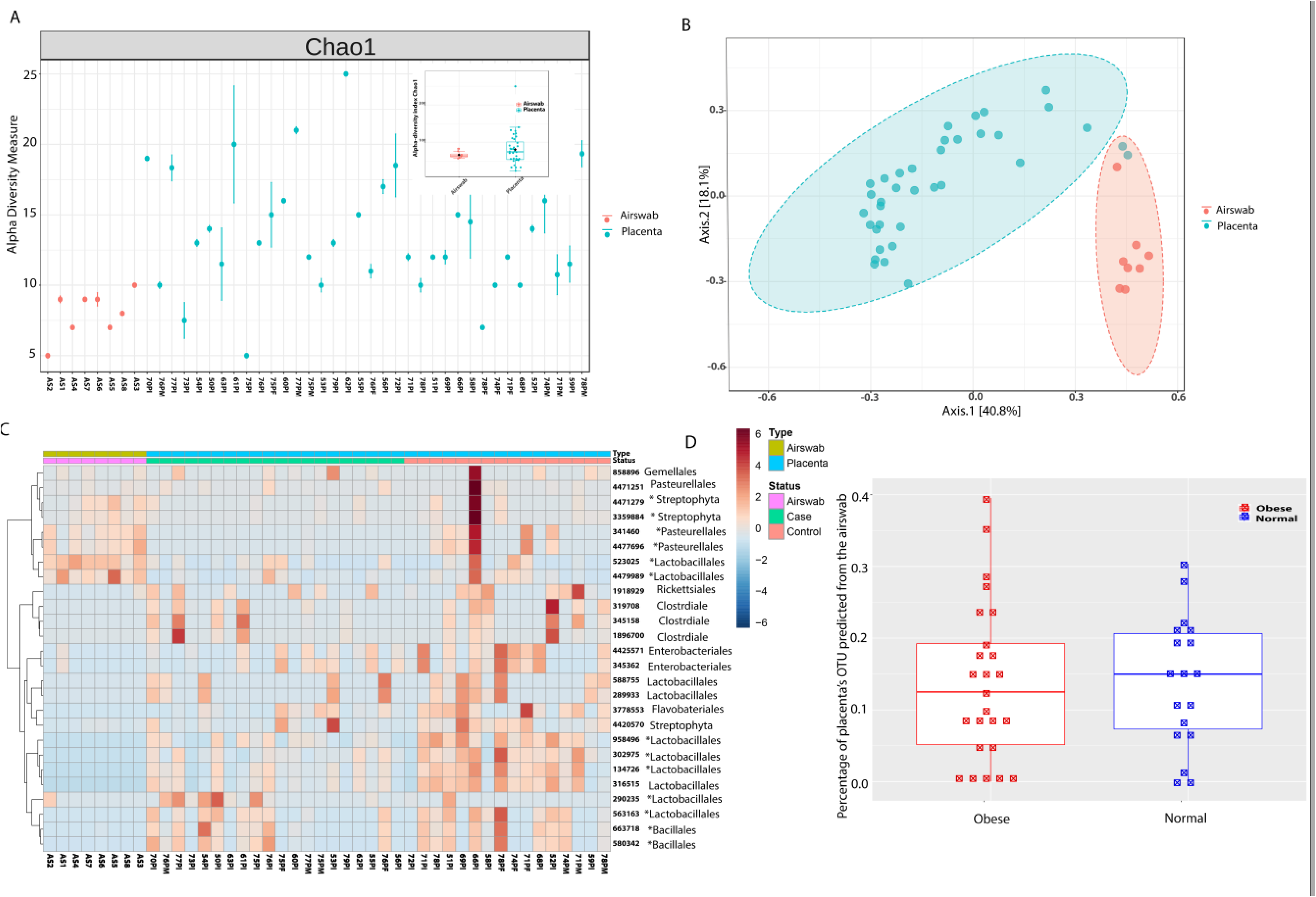
(A) Alpha diversity –Choa1 metric among samples. Boxplot summarizes the OTU diversity difference in placentas and airswabs (t-test, p=6.694E-07). (B) Principle coordinate analysis (PCoA) plot of placenta and airswab clusters, showing two distinct clusters. (C) Heatmap showing different OTUs in placenta samples (pre-pregnant obese and normal pre-pregnant weighted groups) and airswabs.

We further illustrate the OTUs across samples in a heatmap (Figure 2C). Consistent with the difference observed at the summary statistics level, placenta samples and airswabs show clear difference in the types of bacterial OTUs. In particular, 6 OTUs were more abundant in air swabs compared to placenta: *Pasteurellales* (Greengenes OTU ID 4477696, Geengenes OTU ID 341460), *Lactobacillales* (Geengenes OTU ID 523025, Greengenes OTU ID 4479989), and *Streptophyta* (Geengenes OTU ID 3359884, Greengenes OTU ID 4471279). On the contrary, placenta samples contained 13 OTUs that are more abundant compared to the airswabs: *Enterobacteriales* (Greengenes OTU ID 4425571, Greengenes OTU ID 345362), *Lactobacillales* (Greengenes OTU ID 289933, Greengenes OTU ID 588755, Greengenes OTU ID 958496, Greengenes OTU ID 302975, Greengenes OTU ID 134726, Greengenes OTU ID 316515, Greengenes OTU ID 290235, Greengenes OTU ID 563163), *Flavobacteriales* (Greengenes OTU ID 3778553), and *Bacillales* (Greengenes OTU ID 663718, Greengenes OTU ID 580342). We observed 14 significantly different OTUs with (Mann-Whitney U test, FDR < 0.05) (labelled by * in the heatmap), and plotted them in a series of boxplots emphasizing the original count and log-transformed count (Figure S2). To determine the source of taxa in the placenta and how much is attributed from contaminations from the environment, we used the bioinformatics package SourceTracker (20). SourceTracker analysis reported that a median of 14% (min: 0; max: 40%) of the OTUs present in the placental samples could be explained by the bacterial sources from the air contamination (Figure 2D), indicating that the majority of the bacteria in placenta are not due to air contamination.

Next, we visualized the taxonomic composition of community through direct quantitative comparison of relative abundance (Figure 3). As shown by the stacked bar chart of bacterial taxa in Figure 3A and 3B, placenta samples have much higher abundance of bacteria as compared to airswabs, at the genus level. Moreover, the placenta samples contain distinct microbiome populations from those in airswabs (Figure 3B). In particular, *Lysinibacillus* and *Lactobacillus* are much more abundant in placenta, whereas *Haemophilus* and *Streptoccoccus* are much less prevalent in placenta samples. Additionally, *Chryseobacterium* and *Enterococcus* are only present in placentas but not in airswabs. They are commensal non-pathogenic bacterial lineages from *Bacteroidetes* phylum. This observation is in accordance with other earlier reports (2, 5).

**Figure 3:**
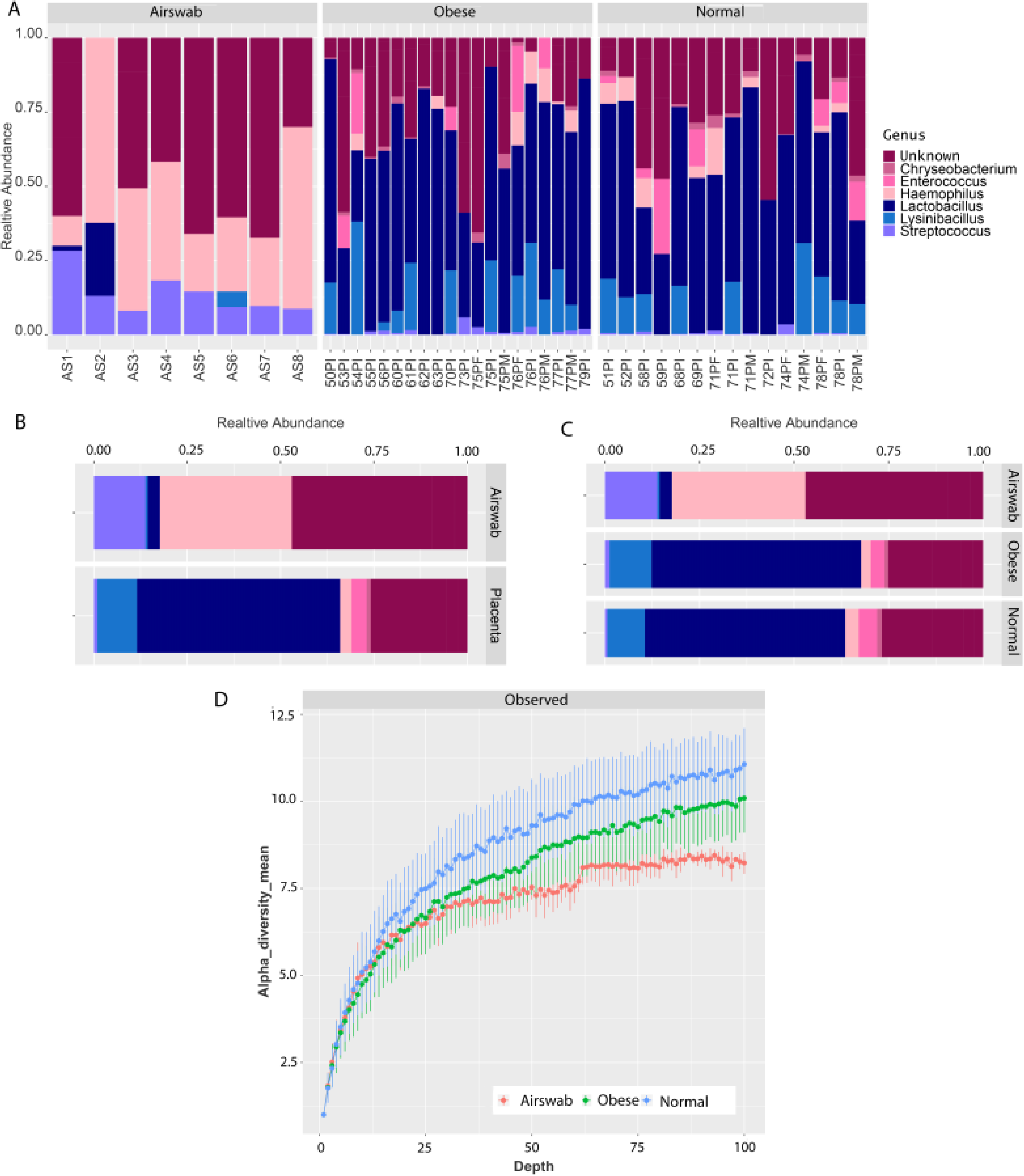
(A) The community structure of all placenta samples at genus level. (B) Relative abundance of bacteria in placenta vs airswab samples. (C) Relative abundance of bacteria grouped by case, control and airswab. (D) The rarefaction curves of airswab (red), pre-pregnant obese (green) and normal pre-pregnant weight samples (blue), at different sequencing depths (x-axes) versus observed alpha diversity (y axes).

### Placental microbiomes of pre-pregnant obese mothers are less diverse than those of mothers of normal pre-pregnant weights

The heatmap of OTUs shows that the placentas of pre-pregnant obese mothers have both less bacterial abundance and diversity, compared to mothers of normal pre-pregnant weights (Figure 2C). It is worth noticing that control sample 66PI shows particularly high bacterial biomass compared to other control samples, possibly indicating an infection. We thus excluded this sample from the following comparisons between cases and controls.

Confirming the observation through OTUs, the average relative abundance of *Lactobacillus* (Mann-Whitney U test, p value =0.01) is significantly lower in obese samples, compared to normal weight samples (Figure 3C), even though there are significant variations among individuals. Additionally, *Haemophilus* has less relative percentage in the obese group, however the difference is not significant (p value =0.24). Previously *Haemophilus* was observed less abundant in the saliva microbiome of obese subjects, compared to those normal controls(21). The overall species richness, measured by alpha-diversity – Chao1 metric, in the rarefaction curve (Figure 3D), is less in pre-pregnant obese samples compared to control samples (t-test, p value = 6.53E-05), across all read depths.

### Placental microbiomes are different from the maternal to fetal side

The placenta samples were collected from three different regions of the placenta: maternal side, intermediate layer and fetal side. We investigated the microbiome abundance and compositions among these three regions (Figure 4). All three placenta regions share most genus types (Figure 4B). Among them, *Lactobacillus*, the dominant taxa in all three layers, shows decreasing relative percentages from the maternal to fetal side (Figure 4B). Additionally, the overall richness (measured by alpha-diversity) is lower in the fetal side, compared to the maternal (p value =0.01) and intermediate layer (p value =0.03), as shown in the rarefaction curve (Figure 4C). Collectively, there appear trends of decrease in both bacterial diversity and the frequency of *Lactobacillus*, from the maternal to the fetal side.

**Figure 4:**
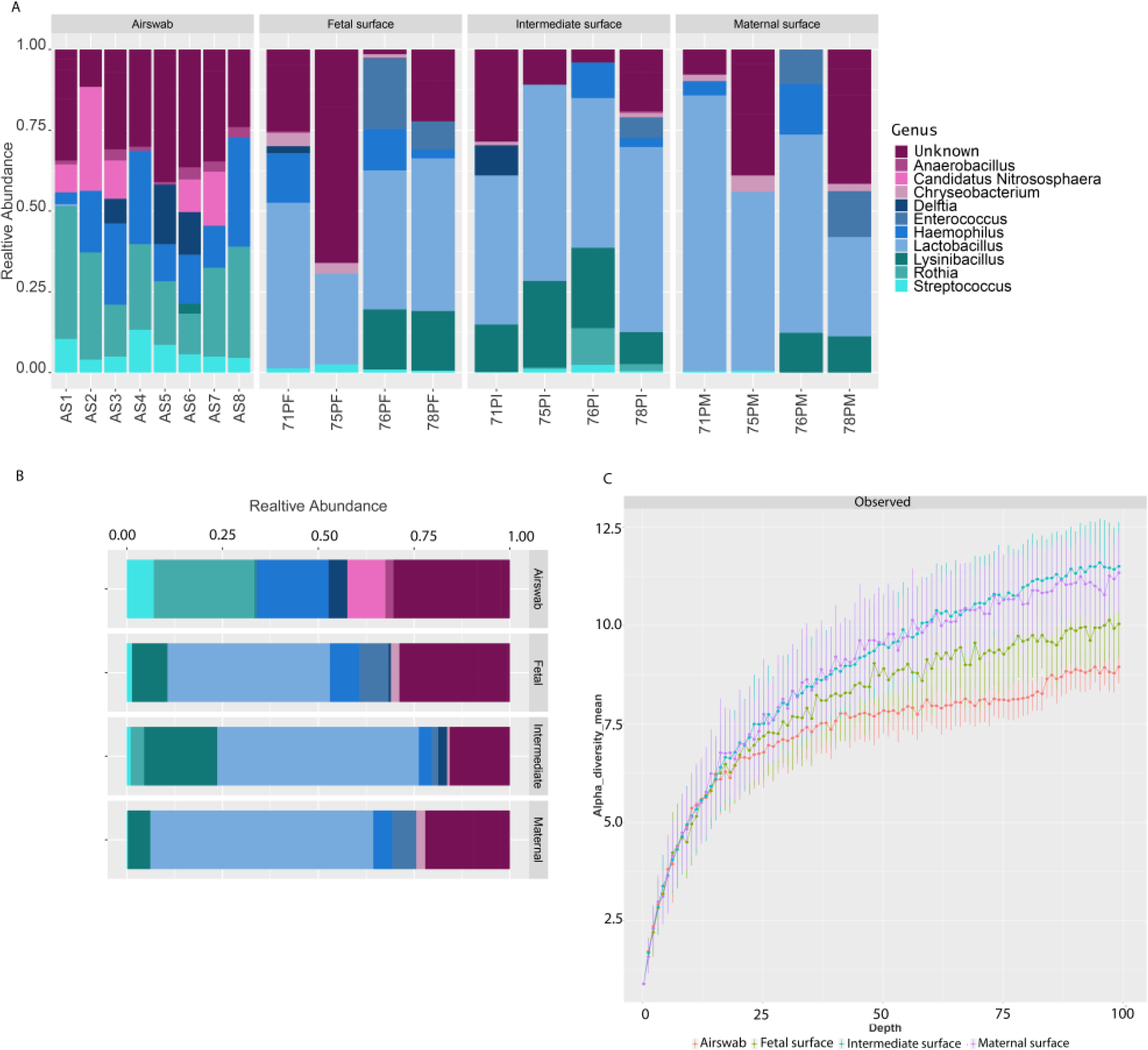
(A) Community structure of placenta samples at genus level obtained from three different placenta layers (maternal surface, intermediate layer, and fetal surface). (B) Relative abundance of placenta microbiome at different placenta surfaces. (C) The rarefaction curves of airswab (red), fetal side (green), intermediate layer (blue), and maternal side (purple), at different sequencing depths (x-axes) versus the alpha diversity (y axes).

## Discussion

In this study, we sought to characterize the biogeography of the placental microbiome in obese and non-obese mothers by performing targeted 16S sequencing of the V4 hypervariable region using an optimized protocol to enrich low bacterial biomass. We applied this protocol to a placenta maternal obesity cohort of women going through elective C-sections, with stringent controls for possible environmental contaminations. We not only confirmed the existence of a unique placenta microbiome distinct from environmental controls, but also found that pre-pregnant obese mothers have reduced bacterial diversity overall. Moreover, within the same placenta, the overall diversity of bacteria appear to decrease from the maternal to fetal side.

We included several careful controls for possible environmental contaminants. We selected women undergoing C-section, rather than those giving birth vaginally, in order to avoid bacterial contaminations from the vaginal region and the nonsterile delivery room. We also used airswabs from the delivery room and laboratory, as well as samples of unopened lab reagents. Our results show strong evidence that taxa contained placental samples are distinct from those in airswab samples, ruling out that the microbiome in placenta is mostly due to contaminations during the experimental procedures. Placentas uniquely have commensal bacteria including *Enterococcus, Lactobacillus* and *Chryseobacterium*, whereas the airswab samples have largely airway-associated taxa (eg. *Haemophilus* and *streptococcus*)(22). *Enterococcus* are Proteobacteria, gram-negative symbionts usually found in the gut. *Lactobacillus* are Firmicutes, gram positive bacteria, and also found in the digestive system where they convert sugar to lactic acid. It was postulated that *Lactobacillius* could transfer from maternal gut to placenta, though the mechanism is unclear (23). *Chryseobacterium*, from the Bacteroidetes phylum, is a type of gram-negative bacteria typically found in milk (24).

Bacterial placenta microbiome diversity is lower in pre-pregnant obese mothers when compared to those non-obese mothers, consistent with previous findings associating obesity with lower microbiome diversity. It was found that oral microbiome were less abundant in obese subjects, compared to normal weighted controls (21); among the mothers who gave spontaneous preterm births, excess gestational weight gains, rather than obesity, were associated with decreased richness in placenta microbiome (25). In our cohort, by experimental design the mothers differed by pre-pregnant BMIs but not by net weight gain, allowing us to directly pinpoint the association between pre-pregnant BMI and microbiome. Additionally, our study only includes full-term births, excluding potential confounding from unknown pathological reasons (such as bacterial infections) which may have existed in the microbiome study in the other pre-term birth cohort (25).

Another interesting finding is the pattern decreasing diversity from the maternal to fetal side in the same placenta. Among them, *Lactobacillus*, the dominant genus in all three layers, also has decreased prevalence from the maternal to fetal side. Interestingly, *Rothia* species are completely absent at the fetal and maternal side, but present at the intermediate layers. Our spatial analysis has suggested the complex bacteria-placenta interactions dependent on localization, and possibly with the fetus in utero.

## Conclusion

Using careful controls for environmental contamination and an enrichment protocol optimized for low bacterial biomass samples, we have confirmed that a unique placenta microbiome does exist. The placental microbiome of pre-pregnant obese weighted mothers is less diverse compared to the normal weighted mothers. Lastly, the microbiome is less diverse on the fetal side compared to the maternal and intermediate layers, suggesting differential section for certain bacterial species according to placenta biogeography.

## Author Contributions

LXG envisioned the project, obtained funding, designed and supervised the project and data analysis. RJS, IYC collected the samples. PAB, FMA, TKW and CD carried out the experiments and analysed the data. All authors have read, edited, revised and approved the manuscript.

### Acknowledgements

Dr. Lana X Garmire’s research is supported by grants K01ES025434 awarded by NIEHS through funds provided by the trans-NIH Big Data to Knowledge (BD2K) initiative (http://datascience.nih.gov/bd2k), P20 COBRE GM103457 awarded by NIH/NIGMS, R01 LM012373 awarded by NLM, and R01 HD084633 awarded by NICHD to LX Garmire. Funding was also provided in part by the Department of Obstetrics and Gynecology, University of Hawaii. 16S sequencing was performed by The University of Minnesota Genomics Center.

## Conflict of Interest

The authors disclose no conflict of interest exists.

---

**Supplementary Figure 1:**
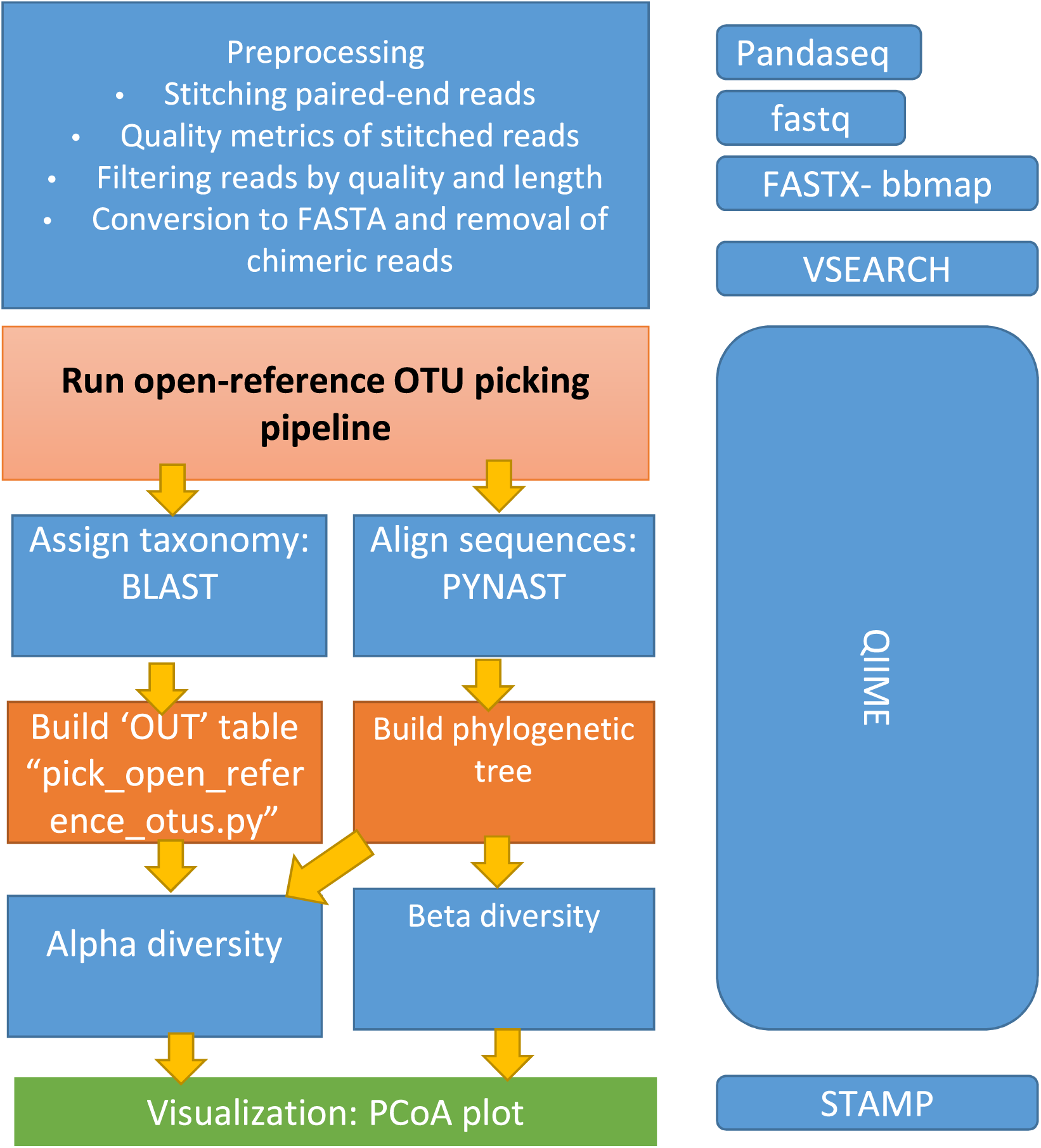
Detailed bioinformatics pipeline used to analyze 16S reads.

**Supplementary Figure 2:**
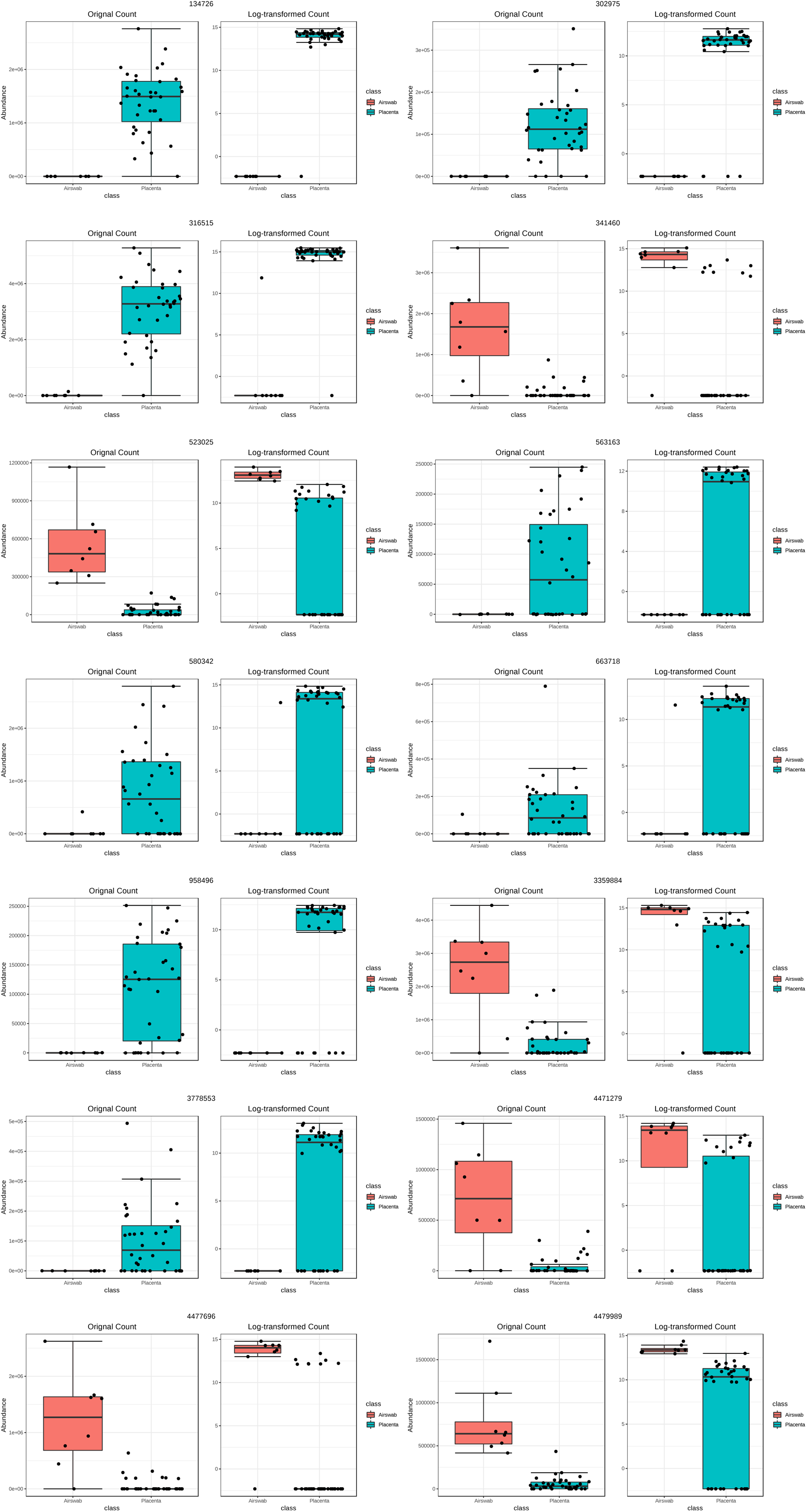
OTUs which are significantly different between placenta and airswab samples (FDR < 0.05)

## References

1. Li X, Watanabe K, Kimura I. 2017. Gut Microbiota Dysbiosis Drives and Implies Novel Therapeutic Strategies for Diabetes Mellitus and Related Metabolic Diseases. Front Immunol 8:1882.

2. Antony KM, Ma J, Mitchell KB, Racusin DA, Versalovic J, Aagaard K. 2015. The preterm placental microbiome varies in association with excess maternal gestational weight gain. Am J Obstet Gynecol 212:653 e1–16.

3. Prince AL, Ma J, Kannan PS, Alvarez M, Gisslen T, Harris RA, Sweeney EL, Knox CL, Lambers DS, Jobe AH, Chougnet CA, Kallapur SG, Aagaard KM. 2016. The placental membrane microbiome is altered among subjects with spontaneous preterm birth with and without chorioamnionitis. Am J Obstet Gynecol 214:627 e1–627 e16.

4. Doyle RM, Alber DG, Jones HE, Harris K, Fitzgerald F, Peebles D, Klein N. 2014. Term and preterm labour are associated with distinct microbial community structures in placental membranes which are independent of mode of delivery. Placenta 35:1099–101.

5. Aagaard K, Ma J, Antony KM, Ganu R, Petrosino J, Versalovic J. 2014. The placenta harbors a unique microbiome. Sci Transl Med 6:237ra65.

6. Lauder AP, Roche AM, Sherrill-Mix S, Bailey A, Laughlin AL, Bittinger K, Leite R, Elovitz MA, Parry S, Bushman FD. 2016. Comparison of placenta samples with contamination controls does not provide evidence for a distinct placenta microbiota. Microbiome 4:29.

7. Zheng J, Xiao X, Zhang Q, Mao L, Yu M, Xu J. 2015. The Placental Microbiome Varies in Association with Low Birth Weight in Full-Term Neonates. Nutrients 7:6924–37.

8. Chu DM, Ma J, Prince AL, Antony KM, Seferovic MD, Aagaard KM. 2017. Maturation of the infant microbiome community structure and function across multiple body sites and in relation to mode of delivery. Nature Medicine 23:314.

9. Jones HE, Harris KA, Azizia M, Bank L, Carpenter B, Hartley JC, Klein N, Peebles D. 2009. Differing prevalence and diversity of bacterial species in fetal membranes from very preterm and term labor. PLoS One 4:e8205.

10. Leiby JS, McCormick K, Sherrill-Mix S, Clarke EL, Kessler LR, Taylor LJ, Hofstaedter CE, Roche AM, Mattei LM, Bittinger K, Elovitz MA, Leite R, Parry S, Bushman FD. 2018. Lack of detection of a human placenta microbiome in samples from preterm and term deliveries. Microbiome 6:196.

11. Parnell LA, Briggs CM, Cao B, Delannoy-Bruno O, Schrieffer AE, Mysorekar IU. 2017. Microbial communities in placentas from term normal pregnancy exhibit spatially variable profiles. Sci Rep 7:11200.

12. Masella AP, Bartram AK, Truszkowski JM, Brown DG, Neufeld JD. 2012. PANDAseq: paired-end assembler for illumina sequences. BMC Bioinformatics 13:31.

13. Edgar RC. 2010. Search and clustering orders of magnitude faster than BLAST. Bioinformatics 26:2460–1.

14. Caporaso JG, Bittinger K, Bushman FD, DeSantis TZ, Andersen GL, Knight R. 2010. PyNAST: a flexible tool for aligning sequences to a template alignment. Bioinformatics 26:266–7.

15. Price MN, Dehal PS, Arkin AP. 2010. FastTree 2--approximately maximum-likelihood trees for large alignments. PLoS One 5:e9490.

16. Caporaso JG, Kuczynski J, Stombaugh J, Bittinger K, Bushman FD, Costello EK, Fierer N, Pena AG, Goodrich JK, Gordon JI, Huttley GA, Kelley ST, Knights D, Koenig JE, Ley RE, Lozupone CA, McDonald D, Muegge BD, Pirrung M, Reeder J, Sevinsky JR, Turnbaugh PJ, Walters WA, Widmann J, Yatsunenko T, Zaneveld J, Knight R. 2010. QIIME allows analysis of high-throughput community sequencing data. Nat Methods 7:335–6.

17. Dhariwal A, Chong J, Habib S, King IL, Agellon LB, Xia J. 2017. MicrobiomeAnalyst: a web-based tool for comprehensive statistical, visual and meta-analysis of microbiome data. Nucleic Acids Res 45:W180–W188.

18. DeSantis TZ, Hugenholtz P, Larsen N, Rojas M, Brodie EL, Keller K, Huber T, Dalevi D, Hu P, Andersen GL. 2006. Greengenes, a chimera-checked 16S rRNA gene database and workbench compatible with ARB. Appl Environ Microbiol 72:5069–72.

19. McMurdie PJ, Holmes S. 2013. phyloseq: An R Package for Reproducible Interactive Analysis and Graphics of Microbiome Census Data. PlOS ONE 8:e61217.

20. Knights D, Kuczynski J, Charlson ES, Zaneveld J, Mozer MC, Collman RG, Bushman FD, Knight R, Kelley ST. 2011. Bayesian community-wide culture-independent microbial source tracking. Nature methods 8:761–763.

21. Wu Y, Chi X, Zhang Q, Chen F, Deng X. 2018. Characterization of the salivary microbiome in people with obesity. PeerJ 6:e4458–e4458.

22. Rogers GB, Carroll MP, Zain NM, Bruce KD, Lock K, Walker W, Jones G, Daniels TW, Lucas JS. 2013. Complexity, temporal stability, and clinical correlates of airway bacterial community composition in primary ciliary dyskinesia. J Clin Microbiol 51:4029–35.

23. Walker RW, Clemente JC, Peter I, Loos RJF. 2017. The prenatal gut microbiome: are we colonized with bacteria in utero? Pediatric Obesity 12:3–17.

24. Ravel J, Gajer P, Abdo Z, Schneider GM, Koenig SS, McCulle SL, Karlebach S, Gorle R, Russell J, Tacket CO, Brotman RM, Davis CC, Ault K, Peralta L, Forney LJ. 2011. Vaginal microbiome of reproductive-age women. Proc Natl Acad Sci U S A 108 Suppl 1:4680–7.

25. Antony KM, Ma J, Mitchell KB, Racusin DA, Versalovic J, Aagaard K. 2015. The preterm placental microbiome varies in association with excess maternal gestational weight gain. American journal of obstetrics and gynecology 212:653.e1-653.16.

